# A Hydroponic-Based Bioassay to Facilitate *Plasmodiophora brassicae* Phenotyping

**DOI:** 10.1101/2023.05.13.540618

**Authors:** Rasha Salih, Anne-Sophie Brochu, Caroline Labbé, Stephen E. Strelkov, Coreen Franke, Richard Bélanger, Edel Pérez-López

## Abstract

Clubroot, caused by the obligate parasite *Plasmodiophora brassicae*, is one of the most devastating diseases affecting the canola/oilseed rape (*Brassica napus*) industry worldwide. Currently, the planting of clubroot-resistant (CR) cultivars is the most effective strategy used to restrict the spread and the economic losses linked to the disease. However, virulent *P. brassicae* isolates have been able to infect many of the currently available CR cultivars, and the options to manage the disease are becoming limited. Another challenge has been achieving consistency in evaluating host reactions to *P. brassicae* infection, with most bioassays conducted in soil and/or potting medium, which requires significant space and can be labour intensive. Visual scoring of clubroot symptom development can also be influenced by user bias. Here, we have developed a hydroponic bioassay using well-characterized *P. brassicae* single-spore isolates representative of clubroot virulence in Canada, as well as field isolates from three Canadian provinces, in combination with canola inbred homozygous lines carrying resistance genetics representative of CR cultivars available to growers in Canada. To improve the efficiency and consistency of disease assessment, symptom severity scores were compared with clubroot evaluations based on the scanned root area. According to the results, this bioassay offers a reliable, less expensive, and reproducible option to evaluate *P. brassicae* virulence, as well as a means to identify which canola resistance profile(s) may be effective against particular isolates. This bioassay will contribute to the breeding of new CR canola cultivars and the identification of virulence genes in *P. brassicae* that could trigger resistance and have been very elusive to this day.

## INTRODUCTION

Canola (*Brassica napus* L.) was established as a crop in Canada 50 years ago, resulting in a thriving industry with a wide international reach. The industry has grown significantly, and each year 20 million tons of canola are produced by almost 43,000 farmers across Canada, representing close to $30 billion to the Canadian economy (CCGA 2022). Canola is predominantly produced in the Prairie Provinces of Manitoba, Alberta, and Saskatchewan, and to some extent in Quebec, British Columbia, and Ontario (Strelkov et al. 2014). Because of this intensified production, canola has become the target of many diseases affecting yield and the stability of the industry. Among them, clubroot disease caused by the intracellular obligate protist *Plasmodiophora brassicae* (Wor.) is a devastating disease leading to annual yield losses estimated at over 20% of the annual production (Javed et al. 2022).

Clubroot is characterized by the formation of galls or clubs in the roots of susceptible cruciferous crops as the result of cellular hyperplasia and hypertrophy (Javed et al. 2022). Aboveground symptoms include wilting, yellowing, stunting, desiccation and eventually the death of the plant (Kageyama and Asano 2009; Javed et al. 2022). During its life cycle, the clubroot pathogen produces resting spores that can survive up to 15 years in the soil (Kageyama and Asano 2009; Javed et al. 2022). The resting spores remain in the soil until a new host is found, where they germinate to penetrate root hairs and start a new cycle that culminates with the formation of the galls and the release of new spores into the soil (Javed et al. 2022).

Since the emergence and identification of *P. brassicae* in the 19^th^ century, many efforts have been undertaken to manage this pathogen and to stop its spread. Measures like crop rotation, the application of chemical products to increase soil pH, and sanitation, although effective in the short term, have not been as effective as the use of clubroot-resistant cultivars (Rahman et al. 2014; Javed et al. 2022). The first clubroot-resistant (CR) canola cultivar became commercially available in 2009. This was soon followed by various other resistant cultivars from different seed companies and these newly introduced varieties exhibited strong resistance to the predominant *P. brassicae* pathotypes (Strelkov et al. 2018).

Currently, there are more than 40 registered clubroot-resistant canola cultivars in the market, making genetic resistance the most popular and effective management option for this disease. Unfortunately, in the arms race between plants and pathogens, there is always a winner, and after years of continuous use of CR canola cultivars, new virulent *P. brassicae* pathotypes have emerged; by the year 2021, more than 6,000 canola fields across Canada were infested by the clubroot pathogen (Javed et al., 2022; Vañó et al. 2023).

To overcome this issue, an improved understanding of the virulence deployed by *P. brassicae* is necessary. Since the 1950s, scientists have been trying to classify *P. brassicae* isolates based on the physiological response of the hosts (Ayers 1957; Williams and Walker 1963). The initial classification as races, was later changed by pathotypes, identifying *P. brassicae* isolates with differential virulence profiles (Somé et al. 1996). Many efforts have been undertaken to develop an accurate and reproducible pathotyping system for *P. brassicae* that can fit needs (Williams 1966; Somé et al. 1996; Strelkov et al. 2018; Pang et al. 2020). Three clubroot pathotyping systems have been widely used by growers, breeders, and researchers: the Williams’ clubroot differential (Williams 1966), the European clubroot differential (ECD) (Buczacki et al. 1975), and Somé clubroot differential (Somé et al. 1996). However, none of those clubroot differential systems included CR cultivars, limiting their applicability. This motivated the development of a new clubroot differential system known as the Canadian Clubroot Differential (CCD) using 13 hosts, including a CR canola (Strelkov et al. 2018). Unfortunately, there are several drawbacks associated with these systems: (*i*) lack of sufficient representation of the CR profiles in commercially available canola, (*ii*) labour-intensive bioassays, (*iii*) time-consuming process, (*iv*) inconsistencies in the evaluation of symptoms severity based on visual assessments, and (*v*) high costs (Williams 1966; Somé et al. 1996; Strelkov et al. 2018; Pang et al. 2020; Tso et al. 2021; Javed et al. 2022). To help to overcome these limitations, we have developed a new hydroponic bioassay, which coupled with a new root surface area scoring scale, can classify *P. brassicae* isolates as virulent or avirulent. In addition to providing a reproducible variable to evaluate *P. brassicae* pathogenicity, this new bioassay shortened the process from six to eight weeks down to five weeks and lowered considerably the input costs.

## MATERIALS AND METHODS

### Plant material

To establish the reliability of a hydroponic bioassay to phenotype *P. brassicae* isolates and to study canola resistance, a set of four inbred homozygous canola lines (IB), obtained from Nutrien Ag Solutions Canada, was used (Table 1). Four of the IB canola lines are clubroot-resistant, with two of them carrying a single CR gene, each from a different source (CRM – *Brassica napus* cv. ‘Mendel’; CR1 – *B. rapa* Chinese cabbage), while the other two carry more than one CR gene (multigenic clubroot resistance or MGCR) (Table 1). These IB lines have been previously phenotyped by our industry partners against five clubroot pathotypes (Table 1). As a susceptible control, we included in each assay the canola cultivar Westar (Table 1).

**Table 1.** Canola germplasm used to phenotype *Plasmodiophora brassicae* isolates.

| Cultivar/Line | Resistance | R gene(s) | R source | Phenotyping* |  |  |  |  |
| --- | --- | --- | --- | --- | --- | --- | --- | --- |
|  |  |  |  | 3H | 5X | 3A | 2B | 3D |
| Westar | No CR | none | no CR | S | S | S | S | S |
| Inbred-1<br>(IB1) | Single-CR gene | CRM | *B. napus* cv.<br>‘Mendel’ | R | S | S | S | S |
| Inbred-2<br>(IB2) | Single-CR gene | CR1 | *B. rapa*<br>Chinese cabbage | R | S | S | S | S |
| Inbred-3<br>(IB3) | Multi-CR genes | MGCR | *B. napus* Rutabaga | R | R | R | R | R |
| Inbred-4<br>(IB4) | Multi-CR genes | MGCR | *B. napus* Rutabaga | R | R | R | R | R |

\* Information provided by industry partners.

### Plasmodiophora brassicae isolates

Thirteen *P. brassicae* isolates were used to validate the hydroponic bioassay (Table 2). Among these, four were single-spore isolates (SSIs) previously characterized and generated from Alberta field isolates and representing a diverse array of virulence patterns (Table 2) (Askarian et al. 2021). Additionally, we included eight uncharacterized field isolates collected by our group or generously donated by industry partners from infected CR canola cultivars in Alberta, Saskatchewan, and Quebec (Table 2). All *P. brassicae* isolates were propagated and maintained on the canola cv. Westar to generate fresh resting spores for each inoculation as described below.

**Table 2.** *Plasmodiophora brassicae* isolates used in this study.

| Isolate | Province* | Type | Pathotype |  |  | Origin |
| --- | --- | --- | --- | --- | --- | --- |
|  |  |  | CCD | Williams | Somé |  |
| Parcelle 3#3 | QC | Field | - | - | - | This study |
| Parcelle 54#5 | QC | Field | - | - | - | This study |
| 1_NAS_3A_Westar | AB | Field | - | - | - | This study |
| 3_NAS_CR breaking_3AB | AB | Field | - | - | - | This study |
| 5_NAS_Rb_1AB | AB | Field | - | - | - | This study |
| SK04 | SK | Field | - | - | - | This study |
| SK16 | SK | Field | - | - | - | This study |
| SK26 | SK | Field | - | - | - | This study |
| 183-14-SS4 | AB | SSI | 4A | 4 | P<sub>1</sub> | Askarian et al. (2020) |
| 187-14-SS4 | AB | SSI | 5X | 5 | P<sub>3</sub> | Askarian et al. (2020) |
| 4-14-SS2 | AB | SSI | 6B | 6 | P<sub>3</sub> | Askarian et al. (2020) |
| 175-14-SS1 | AB | SSI | 6C | 6 | P<sub>3</sub> | Askarian et al. (2020) |

\*QC, Quebec, AB, Alberta, and SK, Saskatchewan

### Inoculum preparation

Resting spores were extracted as previously described with some modifications (Pérez-López et al. 2020). Briefly, clubbed Westar canola roots were washed in 0.25% Tween 20 solution for 5 mins, followed by a 5-mins wash in 70% ethanol and two washes in ddH_2_O for 6 mins. The surface-sterilized clubbed roots were homogenized with 10% (w/v) sucrose solution in a commercial blender. The suspension was passed through eight-layered cheesecloth and the liquid fraction was then centrifuged at 300 rpm for 6 mins to remove root tissue debris. The filtrate was centrifuged at 4500 rpm for 10 mins, the supernatant was carefully discarded, and the pellet was washed twice with ddH_2_O. Spores were resuspended in ddH_2_O and spore density was determined using a hemocytometer adjusting the final spore concentration depending on the assay.

### Hydroponic conditions

For all the hydroponic experiments, canola seeds were germinated in autoclaved vermiculite under controlled conditions, 12 h light/12 h dark, 26°C during the day, 16 °C at night, and 60% humidity. Ten-day-old seedlings were individually transferred into plastic baskets containing hydrotons (Liafor Hydroton, CAD) and carefully placed in a hydroponic system in the greenhouse as previously described with extensive modifications (Lebreton et al. 2018). Each unit in the hydroponic system contained a 10-L container, a 4-cm thick styrofoam sheet in which 30 5-cm holes were drilled to accommodate 5-cm hydroponic baskets/net pots (Fig. 1). Each hydroponic container was filled with 10 L of nutrient solution and aerated with an air stone coupled to an air pump (Marina, CAD) (Fig. 1). An important consideration was to ensure that each seedling’s roots were immersed in the nutrient solution comprised of 4 g of 20-20-20 (Plant-Prod, CAD) and 4.5 g of Epsom salt (MgSO_4_.7H_2_O) at pH 6.5. The nutrient solution was added to the hydroponic tank once a week to keep the volume at 10 L, electric conductivity around 120×10 µs/cm and pH 6.5. Two days post-transplantation, the nutritive solution was inoculated directly with 100 ml of 2 x 10^8^ spores/ml solution, representing approximately 3 ml of inoculum/plant (Fig. 1). Each hydroponic unit fit 30 plants belonging to five different host plants in six replicates (Table 1). The was conducted with a completely randomization, and it was repeated in time including the 13 isolates tested in this study and a unit mock-inoculated with ddH_2_O. All experiments were carried out under greenhouse conditions consisting of temperatures of 23°C to 26°C, 30-50% overall relative humidity, and a 16-h photoperiod (Fig. S1).

**Fig. 1.**
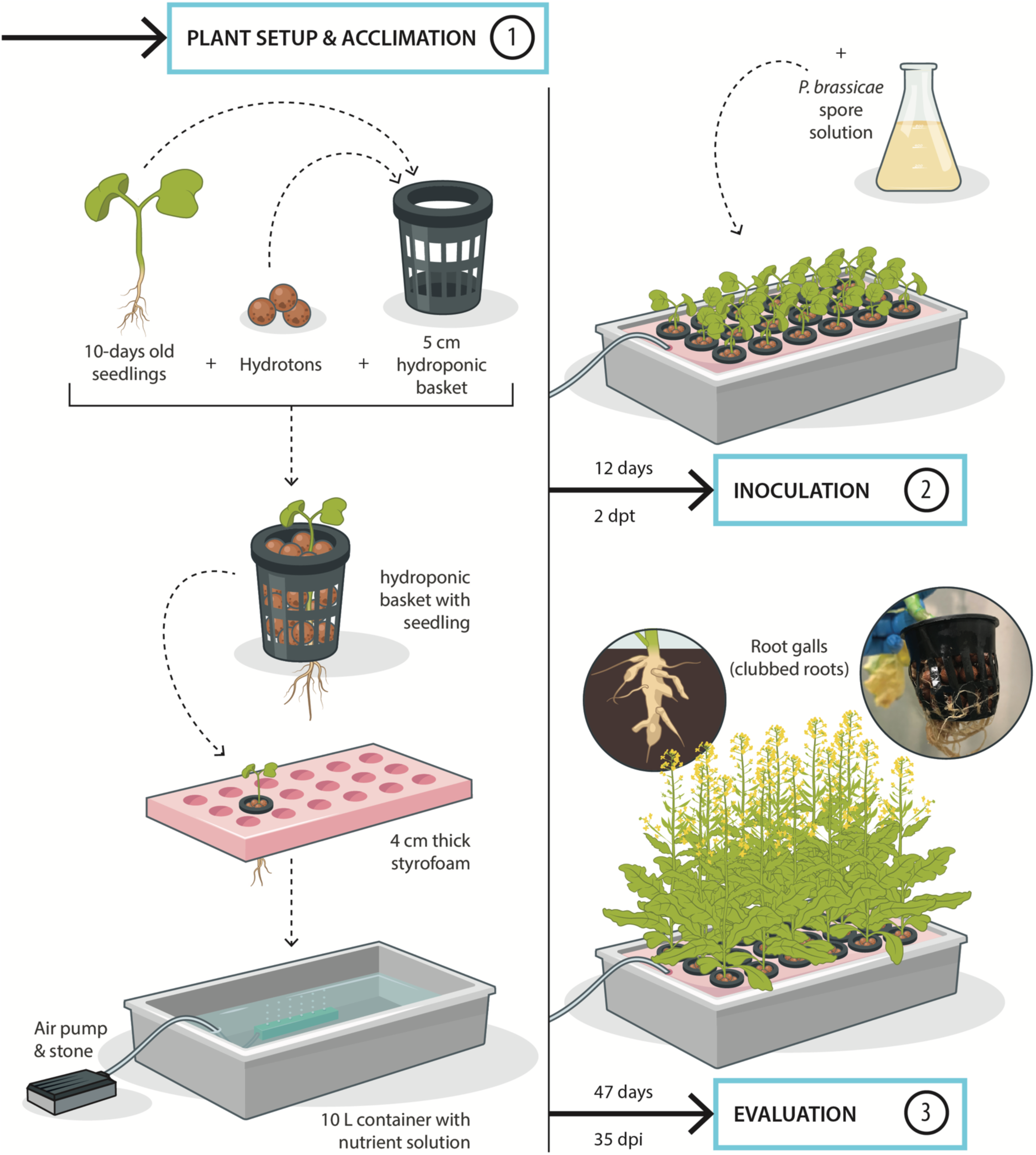
Schematic representation of the hydroponic bioassay developed here to study canola resistance to *Plasmodiophora brassicae*. The process is mainly divided into three parts: (*i*) plant setup and acclimation, (*ii*) inoculation, and (*iii*) evaluation.

### Phenotyping and data analysis

Clubroot symptoms were evaluated 35 days-post inoculations (dpi), recording the number of plants by canola line infected by each isolate evaluated in this study. To rate clubroot symptom development, we used a scale of 0 to 3, where: 0 represents no symptoms/healthy, 1 represents a few small clubs on lateral roots, 2 represents small clubs on the main root and larger clubs on lateral roots, and 3 represents large galls on both main and lateral roots (Kuginuki et al. 1999). In addition, once the symptoms were assessed, all roots were washed and scanned using a WinRHIZO root scanner (Regent Instruments Inc., USA), and the symptoms scale was further validated by comparison with root surface area as a reproducible variable. The statistical significance was determined using a general linear model (GLM). All statistical analyses were performed with R language for statistical computing (R Core Team, 2021).

### Validation of hydroponic bioassay

To determine the reliability of the hydroponic system in comparison with other phenotyping substrate-based assays, we inoculated the four IB canola lines and Westar canola with a field isolate of *P. brassicae* representing pathotype 3A. To test this, the same number of plants was inoculated using the hydroponic bioassay as described above, and in soil/potting medium Veranda mix (Veranda, CAD) (hereafter soil). Seedlings germinated in vermiculite were transplanted to the substrate and inoculated 2-days later with 3 ml of a 2×10^8^ spores/ml inoculum suspension. Symptoms were recorded at 35 dpi, as well as the space occupied by each assay, cost, and other cons and pros associated with hydroponic or substrate-based bioassays.

## RESULTS

### Characterization of *P. brassicae* isolate virulence

Using the hydroponic bioassay, susceptible hosts started to show root swelling at 21 dpi. The symptoms progressed over the next two weeks until the roots developed into the typical galls characteristic of clubroot disease. Westar canola, a susceptible host, displayed a wide range of aboveground symptoms such as stunting, yellowing and death, although the hydroponic solution tended to delay mortality despite a heavily infected root system (Fig. 2). These symptoms were also visible in IB1 and IB2, each carrying a single CR gene and expected to deploy an incompatible (resistant) interaction with at least some of the field isolates tested in this study (Fig. 2). For their part, IB3 and IB4, carrying multiple CR genes, showed incompatible interactions with all the isolates (Fig. 2), and these plants did not show yellowing, stunting or galls on their roots (Fig. 2).

**Fig. 2.**
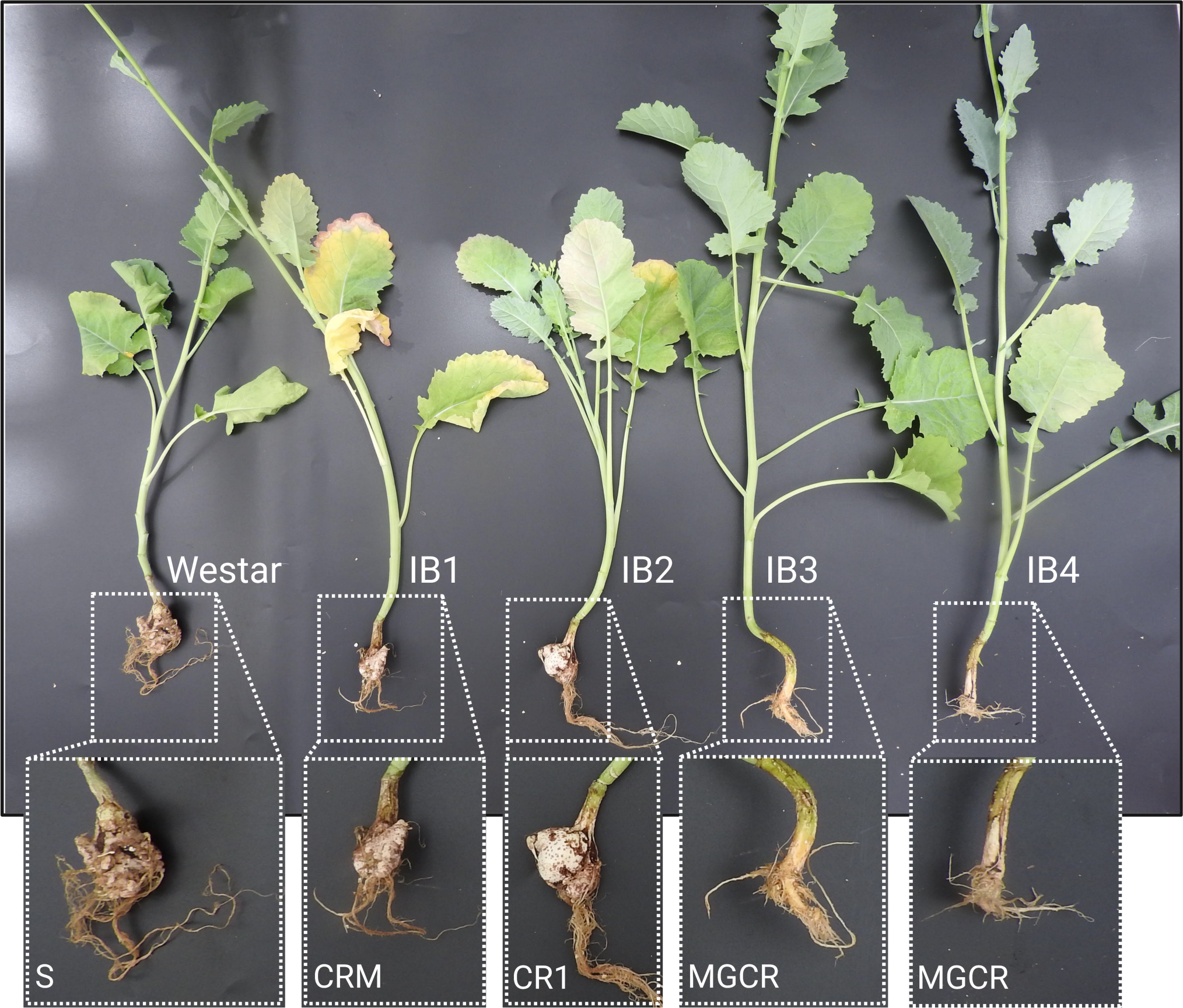
Disease symptoms in five canola hosts used in this study, Westar (susceptible), IB1 (CRM resistance), IB2 (CR1 resistance), IB3 (MGCR resistance), and IB4 (MGCR resistance), infected with *Plasmodiophora brassicae* pathotype 3A using a hydroponic bioassay.

By having compatible and incompatible interactions in the same hydroponic unit, we have confirmed the hydroponic system’s ability to discriminate virulent pathotypes and resistant canola. Alongside Westar canola, IB1 and IB2 were compatible (susceptible) to all the field isolates tested in this study from Alberta, Saskatchewan, and Quebec, with more than 50% of the plants tested consistently infected (Fig. 3). Observations alone identified the 12 isolates tested as virulent to CRM and CR1, but avirulent to the MGCR clubroot resistance profile, also known by the industry as “second-generation” resistance (Fig. 2, Fig. 3, Table S1). Similarly, SSIs were all virulent on the canola lines IB1 and IB2, while an incompatible interaction was observed with IB3 and IB4 carrying multiple CR genes (Fig. 2, Fig. 3, Table S1). Those SSIs were previously pathotyped using the CCD as 4A, 5X, 6B, and 6C, of which, 4A, 5X, and 6B are virulent towards CRM (Mendel-derived resistance), the CR gene present in IB1 (Fig. 2). We found that isolate 175-14-SS1, previously pathotyped as 6C and identified as avirulent on CRM, was virulent on CRM and CR1 in the hydroponic bioassay (Fig. S2).

**Fig. 3.**
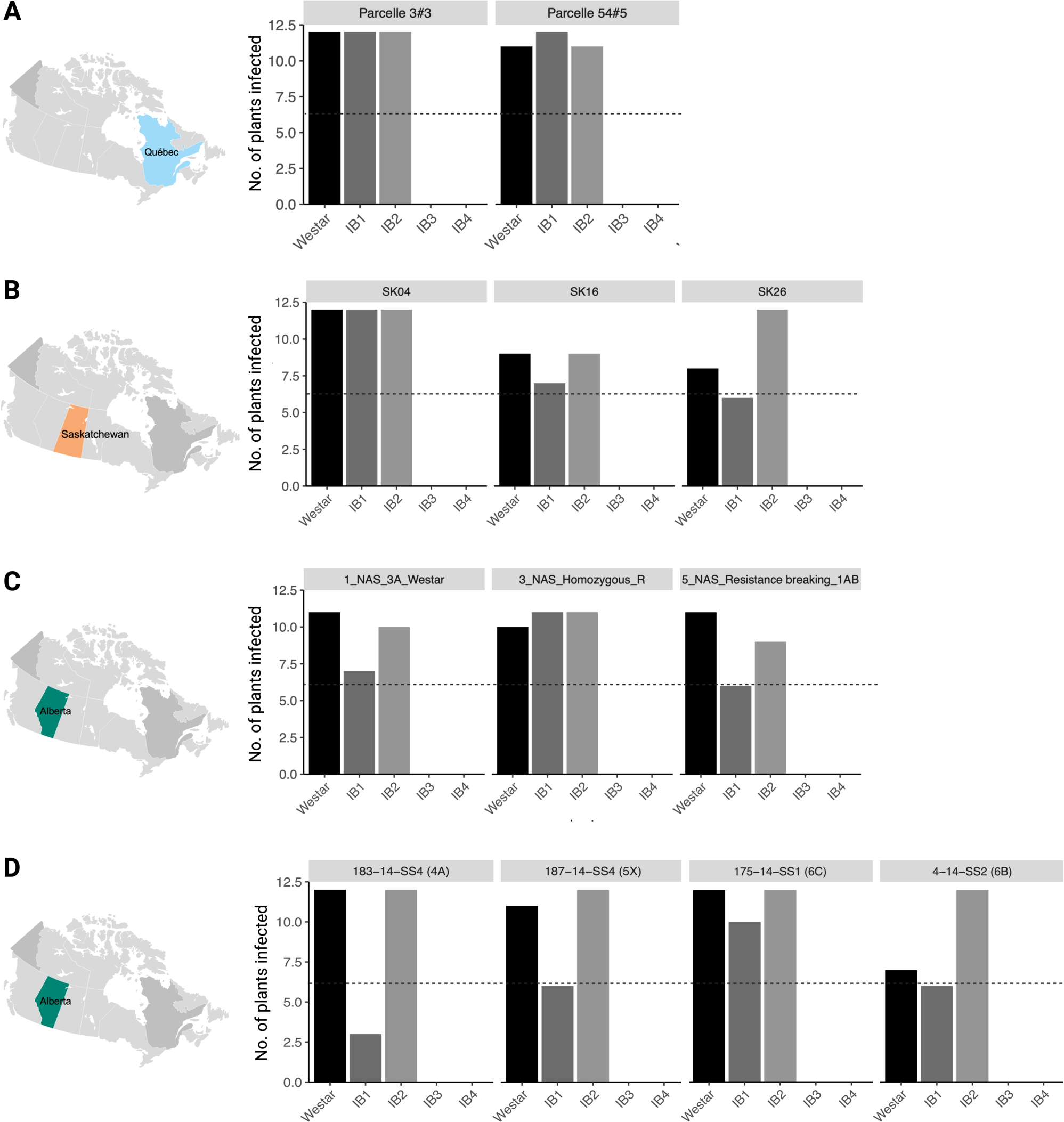
Number of hosts infected by each isolate of *Plasmodiophora brassicae* used in this study. **A**, field isolates from Quebec; **B**, field isolates from Saskatchewan; **C**, field isolates from Alberta; and **D**, single spore isolates from Alberta.

### Root area-based clubroot disease severity score

To facilitate clubroot disease evaluation using the hydroponic bioassay, we applied a clubroot disease severity scale ranging from level 0 to level 3 (Fig. 4A). To validate the use of the root surface area obtained with the WinRHIZO platform (Fig. 4A), we selected 30 plants randomly from each level and compared their root areas; this analysis indicated that the level of infection has a statistically significant effect on the size of the gall and that the root area can be used as a quantitative variable to rate clubroot disease (Fig. 4B, Table S2). The average value of the root area of those plants scored as 0 was 3.9 cm^2^, for those scored as 1 or 2, it was 20.1 cm^2^ or 24.6 cm^2^, respectively, and for those scored as 3, it was 49.5 cm^2^ (Fig. 4B, Table S2). In this analysis, we did not consider the effect of host or isolate, including into each category different combinations (Table S2), and making this variable more robust and flexible for use in different scenarios.

**Fig. 4.**
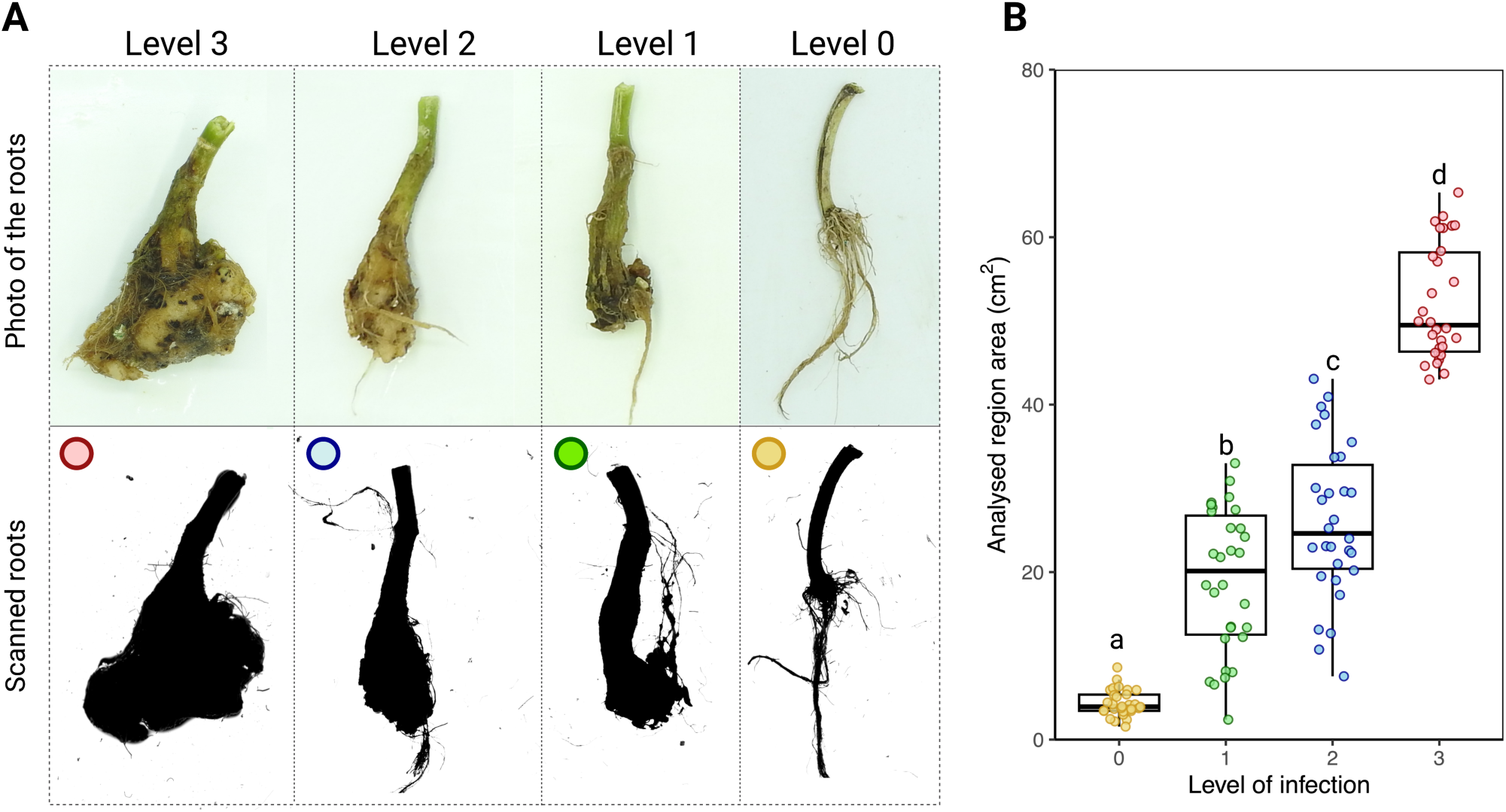
Clubroot disease (*Plasmodiophora brassicae*) severity score to phenotype canola plants using the hydroponic bioassay developed in this study. **A**, example of roots and their respective scan of plants classified into levels 0-3. **B**, comparison of the root area for each level of clubroot disease severity. Letters a-d indicate statistical differences among the levels with *p* < 0.05, n= 30.

### Validation of the hydroponic system

Considering that *P. brassicae* phenotyping is usually carried out in soil, potting medium, or a mixture of both, we wanted to confirm if there was a correlation of infection patterns between systems. Using pathotype 3A, we observed a similar level of infection in both the hydroponic system and soil-based system (Fig. 5). We also observed that Westar, IB1, and IB2 canola were susceptible to *P. brassicae* pathotype 3A with both phenotyping methods, while IB3 and IB4 canola were clubroot resistant, consistent with the information provided by the industry (Fig. 5A-B, Table 1). The similarities were not only in the number of plants infected with each system but also in the severity of infection on each host (Fig. 5B). Another observation made during this comparison was that plants evaluated using hydroponics developed more slowly than in soil (Fig. 5A), contributing to improved space management and the production of less plant debris.

**Fig. 5.**
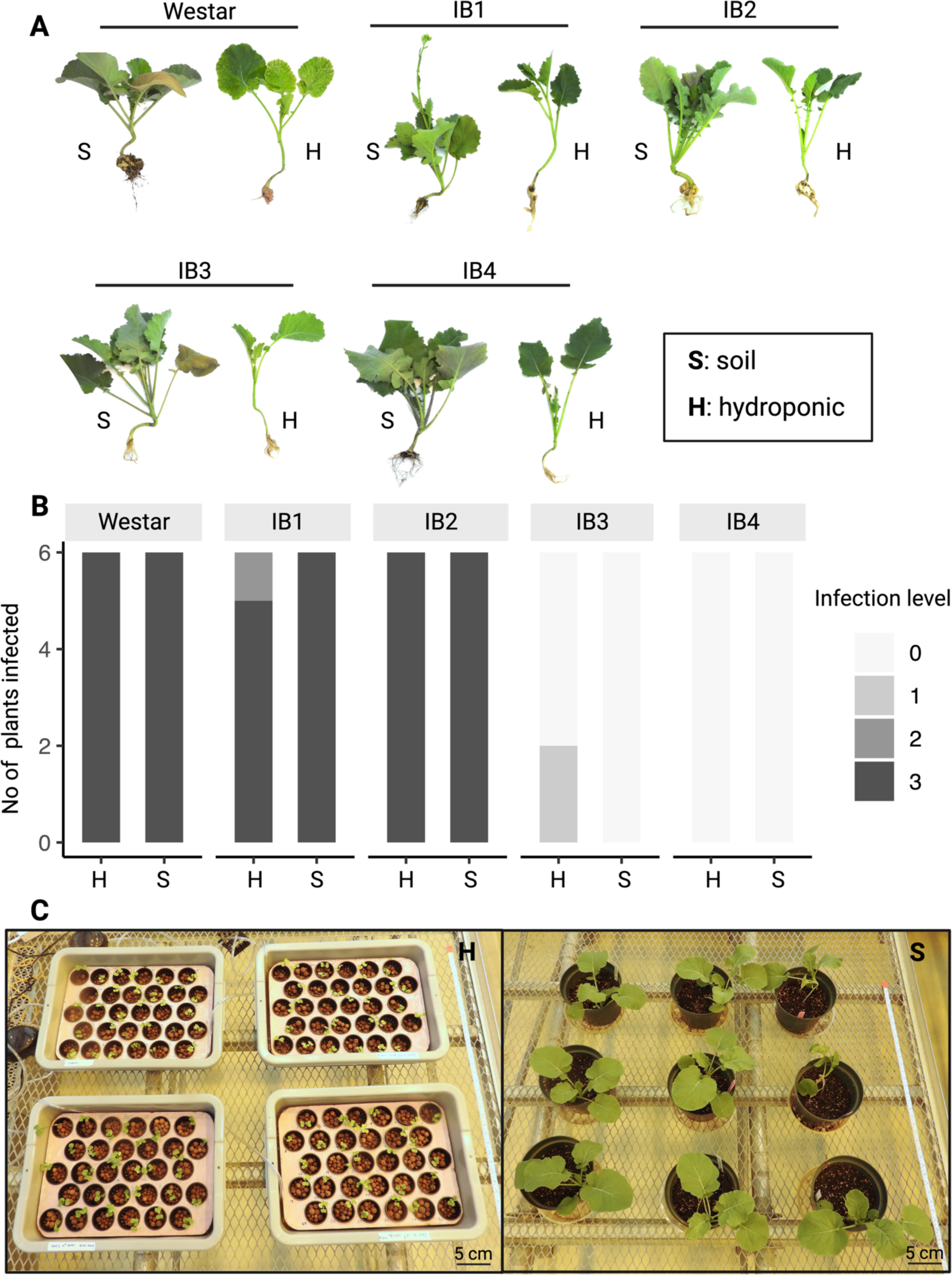
Reliability of the hydroponic system in comparison with soil-based phenotyping to study canola resistance to *Plasmodiophora brassicae*. **A**, phenotypic and developmental variation of the canola cultivars infected using the hydroponic bioassay vs soil-based phenotyping. **B,** level of infection for each cultivar. **C,** space requirements of the hydroponic bioassay vs soil-based phenotyping.

While comparing the hydroponic bioassay with soil inoculation, we also recorded the advantages of this system over substrate-based phenotyping (Table S3). The most notable advantage is how we can fit 120 plants in 1 m^2^, while in the same space, we can fit less than 20 plants using soil or potting medium (Fig. 5C). From a sustainability point of view, the new bioassay requires less water per plant and produces less non-reusable waste (Table S3).

## DISCUSSION

This study describes a hydroponic bioassay that can characterize *P. brassicae* isolates while connecting their virulence to the canola resistance profile. A characteristic of the clubroot pathogen that facilitates the use of a hydroponic system is that following the germination of the resting spores, the next stage of the pathogen life cycle is a biflagellate zoospore (Javed et al. 2022). These zoospores use the aqueous medium in the soil to reach the root hairs, encyst, and penetrate the roots to complete their life cycle (Javed et al. 2022). The question of how good of ‘swimmers’ the clubroot zoospores has not been completely answered (Kageyama and Asano 2009; Javed et al. 2022), but we have confirmed that in the conditions presented here, the infection takes place, and the symptoms are clearly visible more quickly than using soil or potting medium-based phenotyping.

Hydroponic bioassays have been previously used to study clubroot in Chinese cabbage (*Brassica rapa*) (Park et al. 2008), cabbage (*Brassica oleracea*) (Luo et al. 2014), thale cress (*Arabidopsis thaliana*) (Auer 2021), and radish (*Raphanus sativus*) (Li et al. 2022). While those protocols have been partially adapted to study clubroot disease in canola (Chen et al. 2019; Jiang et al. 2022), this study is the first hydroponic bioassay developed to phenotype the clubroot pathogen while studying clubroot resistance in canola. The observation of compatible and incompatible interactions provides evidence that *P. brassicae* can express its effector repertoire to induce the disease in susceptible hosts and trigger plant immunity in those CR canola lines under the conditions described here.

During this first validation, we confirmed which *P. brassicae* isolates are virulent on single CR gene canola in Canada. All field isolates tested using the hydroponic bioassay were able to infect canola lines IB1 and IB2, carrying CRM and CR1, respectively. Those field isolates came from Alberta, Saskatchewan, and Quebec, and all were collected from CR canola varieties from diverse seed companies. In western Canada, the presence of CR-breaking *P. brassicae* isolates has been widely reported (Askarian et al. 2021; Hollman et al. 2021), with recent reports in Ontario (Drury et al. 2021). Here, we report for the first time the presence of CR-breaking isolates in Quebec, the farthest east they have been identified to date and underscoring the threat posed by clubroot to Canadian canola growers.

Several clubroot disease scores have been proposed and used to calculate the disease index (Horiuchi and Hori 1980; Siemens et al., 2002; Salih and Pérez-López, 2021). However, ranking the infected roots via visual estimation is inefficient in terms of labour and costs, and can be subject to operator bias (Salih and Pérez-López, 2021). High-throughput phenotyping using imaging methods has been widely explored to identify disease resistance to root pathogens like the pathogenic oomycetes *Aphanomyces euteiches* infecting peas (Desgroux et al. 2018), the fungal pathogen *Verticillium dahliae* (Bolek et al. 2005), and the oomycete *Pythium ultimatum* in beets (Luterbacher et al. 2005). These methodologies remove the subjective evaluation and can detect differences in the root architecture that visual assessments are unable to capture, an improvement over previous assessments of clubroot disease in Brassica crops.

The proposed hydroponic bioassay overcomes many of the difficulties encountered with other assays relying on soil or substrate inoculation. Previous studies comparing hydroponic and soil phenotyping systems suggest that the growth of the plants in both systems are almost similar and hydroponic conditions require less water and fertilizer compared to soil, making it highly adaptable and more sustainable (Koseki et al. 2011; Verdoliva et al. 2021). The hydroponic system is also feasible in terms of labour, cost, and the number of plants that can be infected in a particular space. As such, using this hydroponic bioassay offers a reliable alternative to evaluate different canola cultivars simultaneously against different *P. brassicae* isolates to phenotype their virulence. Clubroot phenotyping often uses the root-dip method or injects the inoculum into the surrounding soil of the plant at different growth stages to infect the plants with *P. brassicae* resting spores (Johnston 1968; Liu et al. 2022). The efficiency of these traditional methods of inoculation also varies from one host to the other (Yang et al. 2021).

In conclusion, we have developed a reproducible and reliable hydroponic bioassay for the study of *P. brassicae*-canola interaction. Moreover, we have exploited the use of a quantitative variable to phenotype *P. brassicae* isolates virulence and increased reproducibility and inter-laboratory comparisons, improvements needed by the canola industry. This method will facilitate current efforts deployed by the Canola Council of Canada and seed companies to homogenize clubroot resistance labelling and phenotyping, by breeders to achieve more stable and durable clubroot resistance, and by the scientific community to identify new sources of resistance and *P. brassicae* virulence/avirulence factors to improve understanding of the clubroot pathogen.

## ACKNOWLEDGEMENTS

We would like to thank Alberta Canola Producers Commission and Manitoba Canola Growers for funding this research through the Canola Council of Canada’s Canola Agronomic Research Program (CARP), project number 2021.4. We would like to thank Genome Quebec and FRQNT for their support through the Genomic Integration Program, and the NSERC Discovery program for the support. We would like to thank Nutrien Ag Solutions Canada for their support.

---

**Table S1.** *Plasmodiophora brassicae* isolates infection pattern.

| Province | Isolate | Percentage of infected canola lines (%) |  |  |  |  |
| --- | --- | --- | --- | --- | --- | --- |
|  |  | Westar | IB1 | IB2 | IB3 | IB4 |
| QC | Parcelle 3#3 | 100 | 100 | 100 | 0 | 0 |
| QC | Parcelle 54#5 | 92 | 100 | 92 | 0 | 0 |
| AB | 1_NAS_3A_Westar | 92 | 58 | 83 | 0 | 0 |
| AB | 3_NAS_Homozygous_R | 83 | 92 | 92 | 0 | 0 |
| AB | 5_NAS_Rb_1AB | 92 | 50 | 75 | 0 | 0 |
| SK | SK04 | 100 | 100 | 100 | 0 | 0 |
| SK | SK16 | 75 | 58 | 75 | 0 | 0 |
| SK | SK26 | 67 | 50 | 100 | 0 | 0 |
| AB | 183-14-SS4 | 50 | 75 | 100 | 0 | 0 |
| AB | 187-14-SS4 | 92 | 50 | 100 | 0 | 0 |
| AB | 4-14-SS2 | 58 | 50 | 100 | 0 | 0 |
| AB | 175-14-SS1 | 100 | 83 | 100 | 0 | 0 |

**Table S2.** Information used to validate scanned root area as a quantitative variable to calculate clubroot disease index.

| Host | Isolate | Infection Level | Scanned root area (cm<sup>2</sup>) |
| --- | --- | --- | --- |
| IB1 | SK16 | 2 | 17,2742 |
| IB2 | SK16 | 2 | 25,2406 |
| Westar | SK26 | 2 | 13,1327 |
| IB2 | SK26 | 2 | 23,098 |
| IB2 | SK26 | 2 | 29,639 |
| IB2 | SK26 | 2 | 37,6257 |
| IB2 | SK26 | 2 | 24,0038 |
| IB2 | SK26 | 2 | 33,7832 |
| IB2 | 3_NAS_Homo_R | 2 | 22,5798 |
| IB1 | 6C_SSI | 2 | 23,0254 |
| IB1 | 6C_SSI | 2 | 29,4154 |
| IB1 | 6C_SSI | 2 | 20,1992 |
| IB1 | 6C_SSI | 2 | 22,3069 |
| Westar | 6B_SSI | 2 | 39,7596 |
| IB2 | 6B_SSI | 2 | 40,9354 |
| IB2 | 6B_SSI | 2 | 33,7151 |
| IB2 | 6B_SSI | 2 | 43,0838 |
| IB2 | 6B_SSI | 2 | 28,6257 |
| Westar | Parcelle 3#3 | 2 | 12,69 |
| Westar | Parcelle 54#5 | 2 | 26,264 |
| IB1 | Parcelle 54#5 | 2 | 19,0219 |
| IB1 | 3_NAS_Homo_R | 2 | 19,5096 |
| IB2 | 3_NAS_Homo_R | 2 | 29,4935 |
| IB2 | 5_NAS_Resistance breaking_1<br>_AB | 2 | 22,9354 |
| IB3 | 4A_SSI | 2 | 7,5587 |
| IB1 | 5X_SSI | 2 | 35,5238 |
| IB1 | 5X_SSI | 2 | 30,0527 |
| IB3 | 6C_SSI | 2 | 38,7986 |
| IB1 | 6B_SSI | 2 | 20,9909 |
| IB1 | 6B_SSI | 2 | 10,7484 |
| IB2 | SK04 | 3 | 46,1767 |
| Westar | SK16 | 3 | 45,3599 |
| Westar | SK16 | 3 | 49,1225 |
| IB2 | Parcelle 54#5 | 3 | 57,0918 |
| IB1 | 3_NAS_Homo_R | 3 | 44,9331 |
| IB2 | 1_NAS_3A_Westar | 3 | 45,9199 |
| Westar | 4A_SSI | 3 | 58,346 |
| IB2 | 5X_SSI | 3 | 53,3031 |
| IB2 | 5X_SSI | 3 | 65,3341 |
| IB2 | 5X_SSI | 3 | 61,0983 |
| IB2 | 5X_SSI | 3 | 61,8908 |
| IB2 | 6C_SSI | 3 | 61,3683 |
| IB2 | 6C_SSI | 3 | 47,6593 |
| IB2 | 6C_SSI | 3 | 46,7418 |
| IB1 | SK16 | 3 | 48,9644 |
| Westar | Parcelle 3#3 | 3 | 48,3386 |
| IB2 | Parcelle 3#3 | 3 | 49,9499 |
| Westar | Parcelle 54#5 | 3 | 44,6225 |
| IB1 | Parcelle 54#5 | 3 | 46,1902 |
| IB1 | 3_NAS_Homo_R | 3 | 51,1193 |
| IB1 | 5_NAS_Resistance breaking_1<br>_AB | 3 | 57,6928 |
| IB1 | 5_NAS_Resistance breaking_1<br>_AB | 3 | 43,6954 |
| IB1 | 1_NAS_3A_Westar | 3 | 49,8547 |
| Westar | 5X_SSI | 3 | 46,9335 |
| IB2 | 5X_SSI | 3 | 43,0025 |
| IB1 | 6C_SSI | 3 | 61,4205 |
| IB1 | 6C_SSI | 3 | 54,6644 |
| IB1 | 6C_SSI | 3 | 61,1031 |
| IB1 | 6C_SSI | 3 | 62,5031 |
| IB2 | 6C_SSI | 3 | 47,9612 |
| Westar | Control | 0 | 3,4271 |
| Westar | Control | 0 | 4,1497 |
| Westar | Control | 0 | 5,9342 |
| Westar | Control | 0 | 3,6645 |
| Westar | Control | 0 | 2,3774 |
| Westar | Control | 0 | 2,1716 |
| IB1 | Control | 0 | 6,1419 |
| IB1 | Control | 0 | 3,5097 |
| IB1 | Control | 0 | 8,6206 |
| IB1 | Control | 0 | 1,5484 |
| IB1 | Control | 0 | 4,2013 |
| IB1 | Control | 0 | 2,9806 |
| IB2 | Control | 0 | 3,4142 |
| IB2 | Control | 0 | 5,9368 |
| IB2 | Control | 0 | 7,1613 |
| IB2 | Control | 0 | 3,7548 |
| IB2 | Control | 0 | 3,9045 |
| IB2 | Control | 0 | 6,3206 |
| IB3 | Control | 0 | 5,311 |
| IB3 | Control | 0 | 4,229 |
| IB3 | Control | 0 | 5,3935 |
| IB3 | Control | 0 | 5,1097 |
| IB3 | Control | 0 | 3,3284 |
| IB3 | Control | 0 | 3,449 |
| IB4 | Control | 0 | 2,4568 |
| IB4 | Control | 0 | 4,4284 |
| IB4 | Control | 0 | 3,7606 |
| IB4 | Control | 0 | 3,9484 |
| IB4 | Control | 0 | 3,629 |
| IB1 | 4A_SSI | 0 | 5,9251 |
| Westar | Parcelle 3#3 | 1 | 8,0477 |
| Westar | Parcelle 3#3 | 1 | 8,1697 |
| Westar | Parcelle 3#3 | 1 | 6,8981 |
| Westar | Parcelle 3#3 | 1 | 25,2682 |
| Westar | Parcelle 3#3 | 1 | 27,7214 |
| Westar | Parcelle 3#3 | 1 | 21,7887 |
| IB2 | Parcelle 3#3 | 1 | 27,2395 |
| IB2 | Parcelle 3#3 | 1 | 33,0053 |
| IB2 | Parcelle 3#3 | 1 | 28,2977 |
| IB2 | Parcelle 3#3 | 1 | 13,3867 |
| IB1 | Parcelle 54#5 | 1 | 13,4826 |
| IB1 | Parcelle 54#5 | 1 | 12,2501 |
| IB1 | 4A_SSI | 1 | 28,0625 |
| IB1 | 4A_SSI | 1 | 13,3766 |
| IB1 | 4A_SSI | 1 | 12,0774 |
| IB1 | 4A_SSI | 1 | 25,2145 |
| IB1 | 4A_SSI | 1 | 27,4485 |
| IB1 | 4A_SSI | 1 | 24,2187 |
| IB1 | 6B_SSI | 1 | 2,3777 |
| IB1 | 6B_SSI | 1 | 16,2 |
| IB1 | 6B_SSI | 1 | 30,8903 |
| Westar | Parcelle 3#3 | 1 | 18,4427 |
| Westar | Parcelle 3#3 | 1 | 6,57 |
| IB2 | Parcelle 3#3 | 1 | 17,5645 |
| IB2 | Parcelle 3#3 | 1 | 22,558 |
| IB2 | Parcelle 3#3 | 1 | 22,1922 |
| IB1 | Parcelle 54#5 | 1 | 18,4732 |
| Westar | 6C_SSI | 1 | 28,958 |
| Westar | 6C_SSI | 1 | 7,3858 |
| Westar | 6C_SSI | 1 | 22,3083 |

**Table S3.** Feasibility of hydroponics vs soil-based clubroot phenotyping.

| Feature | Hydroponics | Soil |
| --- | --- | --- |
| Number of plants/m<sup>2</sup> | 120 | 9 |
| Watering | N/A | 2-3 times a week |
| Quantity of liquid fertilizer | 60 ml/ unit | 3 ml/ plant |
| Duration | 47 days | 55 days |

**Fig. S1.** Set up of the hydroponic bioassay described in this study. Numbers indicate sequential steps.

**Fig. S2.**
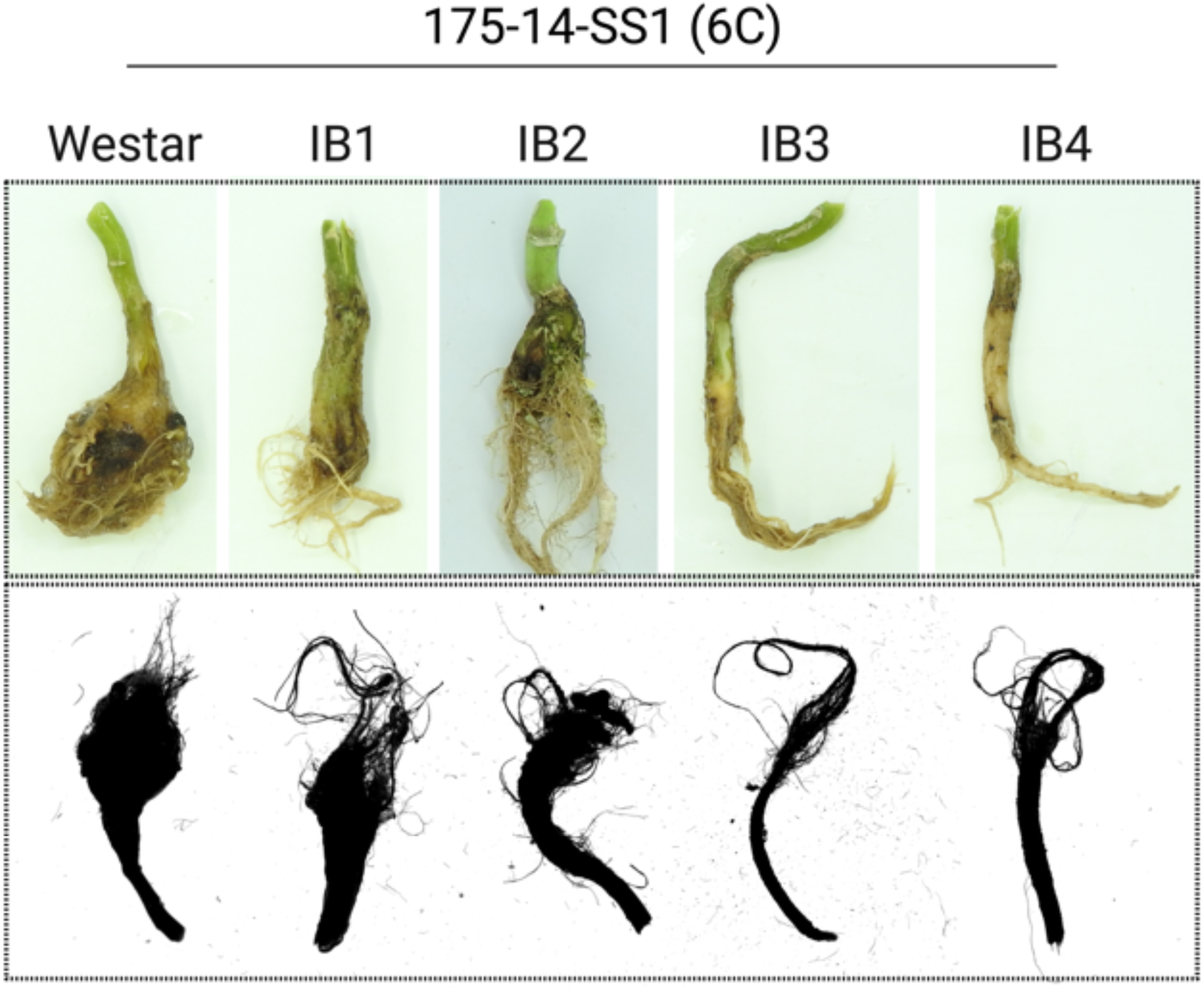
Phenotypes for *Plasmodiophora brassicae* isolate 175-14-SS1, pathotyped as 6C in all canola lines used to validate the hydroponic bioassay.

## REFERENCES

Askarian, H., Akhavan, A., Manolii, V. P., Cao, T., Hwang, S. F. and Strelkov, S. E. 2021. Virulence spectrum of single-spore and field isolates of *Plasmodiophora brassicae* able to overcome resistance in canola (*Brassica napus*). Plant Dis. 105:43–52.

Auer, S. 2021. A costum-made hydroponic culture system to study plant roots during root infection with *Plasmodiophora brassicae*. protocols.io https://dx.doi.org/10.17504/protocols.io.bm3mk8k6

Ayers, G.W. (1957) Races of *Plasmodiophora brassicae*. Can. J. Plant Pathol. 35:923–932.

Barrada, A., Delisle-Houde, M., Nguyen, T. T. A., Tweddell, R. J., and Dorais, M. 2022. Drench Application of Soy Protein Hydrolysates Increases Tomato Plant Fitness, Fruit Yield, and Resistance to a Hemibiotrophic Pathogen. Agronomy. 12:1761.

Bolek, Y., El-Zik, K. M., Pepper, A. E., Bell, A. A., Magill, C. W., Thaxton, P. M., Reddy, O. U. K. 2005. Mapping of verticillium wilt resistance genes in cotton. Plant Science. 168:1581–1590.

Buczacki, S. T., Toxopeus, H., Mattusch, P., Johnston, T. D., Dixon, G. R. and Hobolth, L. A. 1975. Study of physiologic specialization in *Plasmodiophora brassicae*: Proposals for attempted rationalization through an international approach. Trans. Br. Mycol. Soc. 65:295–303.

Canadian canola growers association (CCGA). 2022. Available at: https://www.ccga.ca/policy/Pages/AdvocacyPriorities.aspx. Accessed on 3rd March, 2023.

Chen, W., Li, Y., Yan, R., Xu, L., Ren, L., Liu, F., Zeng, L., Yang, H., Chi, P., Wang, X., and Chen, K. 2019. Identification and characterization of *Plasmodiophora brassicae* primary infection effector candidates that suppress or induce cell death in host and nonhost plants. Phytopathol. 109:1689–1697.

Cortleven, A., Leuendorf, J. E., Frank, M., Pezzetta, D., Bolt, S., and Schmülling, T. 2019. Cytokinin action in response to abiotic and biotic stresses in plants. Plant Cell Environ. 42:998–1018.

Desgroux A., Baudais V. N., Aubert V., Le Roy G., de Larambergue H., Miteul H., et al. 2018. Comparative genome-wide-association mapping identifies common loci controlling root system architecture and resistance to *Aphanomyces euteiches* in pea. Front. Plant Sci. 8:2195.

Drury, S. C., Gossen, B. D., and McDonald, M. R. 2021. Clubroot resistance in canola and brassica vegetable cultivars in Ontario, Canada. Can. J. Plant Sci. 101:730–740.

Hollman, K. B., Hwang, S. F., Manolii, V. P., and Strelkov, S. E. 2021. Pathotypes of *Plasmodiophora brassicae* collected from clubroot-resistant canola *(Brassica napus* L.) cultivars in western Canada in 2017-2018. Can. J. Plant Pathol. 43:622–630.

Javed, M. A., Schwelm, A., Zamani-Noor, N., Salih, R., Silvestre Vañó, M., Wu, J., González García, M., Heick, T. M., Luo, C., Prakash, P., and Pérez-López, E. 2022. The clubroot pathogen *Plasmodiophora brassicae*: A profile update. Mol. Plant Pathol. 24:89–106.

Jiang, X., Su, Y., and Wang, M. 2022. Mapping of a novel clubroot disease resistance locus in Brassica napus and related functional identification. Front. Plant Sci. 13:1014376.

Johnston, T. D. 1968. Clubroot in Brassica: a standard inoculation technique and the specification of races. Plant Pathol. 17:184–184.

Kageyama, K., and Asano, T. 2009. Life cycle of *Plasmodiophora brassicae*. J. Plant Growth Regul. 28:203–211.

Kazan, K., and Lyons, R. 2016. The link between flowering time and stress tolerance. J. Exp. Bot. 67:47–60.

Koseki, S., Mizuno, Y., and Yamamoto, K. 2011. Comparison of two possible routes of pathogen contamination of spinach leaves in a hydroponic cultivation system. J. Food Prot. 74:1536–1542.

Kuginuki, Y., Yoshikawa, H., and Hirai, M. 1999. Variation in Virulence of Plasmodiophora brassicae in Japan Tested with Clubroot-resistant Cultivars of Chinese Cabbage (*Brassica rapa* L. ssp. pekinensis). Eur. J. Plant Pathol. 105:327–332.

Lebreton, A., Labbé, C., De Ronne, M., Xue, A. G., Marchand, G., and Bélanger, R. R. 2018. Development of a simple hydroponic assay to study vertical and horizontal resistance of soybean and pathotypes of *Phytophthora sojae*. Plant Dis. 102:114–123.

Li, J., Huang, T., Lu, J., Xu, X., and Zhang, W. 2022. Metabonomic profiling of clubroot-susceptible and clubroot-resistant radish and the assessment of disease-resistant metabolites. Front. Plant Sci. 13.

Liu, L., Qin, L., Cheng, X., Zhang, Y., Xu, L., Liu, F., Tong, C., Huang, J., Liu, S., and Wei, Y. 2020. Comparing the infection biology of *Plasmodiophora brassicae* in clubroot susceptible and resistant hosts and non-hosts. Front. Microbiol. 11:507036.

Liu, X., Strelkov, S. E., Sun, R., Hwang, S. F., Fredua-Agyeman, R., Li, F., Zhang, S., Li, G., Zhang, S., and Zhang, H. 2022. Histopathology of the *Plasmodiophora brassicae*-Chinese Cabbage Interaction in Hosts Carrying Different Sources of Resistance. Front. Plant Sci. 12:3236.

Lu, T., Ke, M., Lavoie, M., Jin, Y., Fan, X., Zhang, Z., Fu, Z., Sun, L., Gillings, M., Peñuelas, J., and Qian, H. 2018. Rhizosphere microorganisms can influence the timing of plant flowering. Microbiome 6:1–12.

Luo, H. C., Chen, G. K., Liu, C. P., Huang, Y., and Xiao, C. G. 2014. An improved culture solution technique for *Plasmodiophora brassicae* infection and the dynamic infection in the root hair. Australas. Plant Pathol. 43:53–60.

Luterbacher, M. C., Asher, M. J. C., Beyer, W., Mandolino, G., Scholten, O. E., Frese, L., Biancardi, E., Stevanato, P., Mechelke, W., and Slyvchenko, O. 2005. Sources of resistance to diseases of sugar beet in related Beta germplasm: II. Soil-borne diseases. Euphytica. 141:49–63.

Marzougui, A., Ma, Y., Zhang, C., McGee, R. J., Coyne, C. J., Main, D., and Sankaran, S. 2019. Advanced imaging for quantitative evaluation of Aphanomyces root rot resistance in lentil. Front. Plant Sci. 10:383.

Nutrien AG Solutions Canada. 2021. Available at: https://www.nutrienagsolutions.ca/. Accessed on 18th March, 2023.

Pang, W., Liang, Y., Zhan, Z., Li, X., and Piao, Z. 2020. Development of a sinitic clubroot differential set for the pathotype classification of *Plasmodiophora brassicae*. Front. Plant Sci. 1360.

Park, S., Kwak, J., and Yoon, M. 2008. Development of an effective inoculation method for large quantaties of clubroot disease using hydroponics in Chinese cabbage. Korean J. Hortic. Sci. Technol. 26:449–453.

Peng, G., Lahlali, R., Hwang, S. F., Pageau, D., Hynes, R. K., McDonald, M. R., Gossen, B. D., and Strelkov, S. E. 2014. Crop rotation, cultivar resistance, and fungicides/biofungicides for managing clubroot (*Plasmodiophora brassicae*) on canola. Can. J. Plant Pathol. 36:99–112.

R Core Team 2021. R: A language and environment for statistical computing. R Foundation for Statistical Computing, Vienna, Austria. URL https://www.R-project.org/.

Rahman H., Peng G., Yu F., Falk K. C., Kulkarni M., and Selvaraj G. 2014. Genetics and Breeding for Clubroot Resistance in Canadian Spring Canola (*Brassica napus* L.). Can. J. Plant Pathol. 36:122–134.

Rennie, D. C., Holtz, M. D., Turkington, T. K., Leboldus, J. M., Hwang, S. F., Howard, R. J., and Strelkov, S. E. 2015. Movement of *Plasmodiophora brassicae* resting spores in windblown dust. Can. J. Plant Pathol. 37:188–196.

Salih, R., and Pérez-López, E. 2021. Digitalization of Clubroot Disease Index, a Long Overdue Task. Horticulturae. 7:241.

Siemens, J., Nagel, M., Ludwig-Müller, J. & Sacristan, M. D. 2002. The interaction of *Plasmodiophora brassicae* and *Arabidopsis thaliana*: parameters for disease quantification and screening of mutant lines. J. Phytopathol. 150:592–05.

Smith, L. J., and Scarisbrick, D. H. 1990. Reproductive development in oilseed rape (*Brassica napus* cv. Bienvenu). Ann. Bot. 65:205–212.

Somé, A., Manzanares, M. J., Laurens, F., Baron, F., Thomas, G., and Rouxel, F. 1996. Variation for virulence on *Brassica napus* L. amongst *Plasmodiophora brassicae* collections from France and derived single-spore isolates. Plant Pathol. J. 45:432–439.

Strelkov, S. E., and Hwang, S. F. 2014. Clubroot in the Canadian canola crop: 10 years into the outbreak. Can. J. Plant Pathol. 36:27–36.

Strelkov, S. E., Hwang, S. F., Manolii, V. P., Cao, T., Fredua-Agyeman, R., Harding, M. W. et al. 2018. Virulence and pathotype classification of *Plasmodiophora brassicae* populations collected from clubroot resistant canola (*Brassica napus*) in Canada. Can. J. Plant Pathol. 40:284–298.

Tso, H. H., Galindo-González, L., and Strelkov, S. E. 2021. Current and future pathotyping platforms for *Plasmodiophora brassicae* in Canada. Plants 10:1446.

Vañó, M. S., Nourimand, M., MacLean, A., and Pérez-López, E. 2023. Getting to the root of a club– Understanding developmental manipulation by the clubroot pathogen. Semin. Cell Dev. Biol. 149:22–32.

Verdoliva, S. G., Gwyn-Jones, D., Detheridge, A., and Robson, P. 2021. Controlled comparisons between soil and hydroponic systems reveal increased water use efficiency and higher lycopene and β-carotene contents in hydroponically grown tomatoes. Sci. Hortic. 279:109896.

Wang, Y., Koopmann, B., and von Tiedemann, A. 2022. Methods for Assessment of Viability and Germination of *Plasmodiophora brassicae* Resting Spores. Front. Microbiol. 12:4212.

Williams, P. H. 1966. A system for the determination of races of *Plasmodiophora brassicae* that infect cabbage and rutabaga. Phytopathol. 56:624–626.

Williams, P. H., and Walker, J. C. 1963. Races of clubroot in North America. Plant Dis. Rep. 47:608–611.

Yang, H., Yuan, Y., Wei, X., Zhang, X., Wang, H., Song, J., and Li, X. 2021. A new identification method reveals the resistance of an extensive-source radish collection to *Plasmodiophora brassicae* race 4. Agronomy. 11(4):792.

